# Genome-Wide Identification of 5-Methylcytosine Sites in Bacterial Genomes By High-Throughput Sequencing of MspJI Restriction Fragments

**DOI:** 10.1101/2021.02.10.430591

**Authors:** Brian P. Anton, Alexey Fomenkov, Victoria Wu, Richard J. Roberts

## Abstract

Single-molecule Real-Time (SMRT) sequencing can easily identify sites of N6-methyladenine and N4-methylcytosine within DNA sequences, but similar identification of 5-methylcytosine sites is not as straightforward. In prokaryotic DNA, methylation typically occurs within specific sequence contexts, or motifs, that are a property of the methyltransferases that “write” these epigenetic marks. We present here a straightforward, cost-effective alternative to both SMRT and bisulfite sequencing for the determination of prokaryotic 5-methylcytosine methylation motifs. The method, called MFRE-Seq, relies on excision and isolation of fully methylated fragments of predictable size using MspJI-Family Restriction Enzymes (MFREs), which depend on the presence of 5-methylcytosine for cleavage. We demonstrate that MFRE-Seq is compatible with both Illumina and Ion Torrent sequencing platforms and requires only a digestion step and simple column purification of size-selected digest fragments prior to standard library preparation procedures. We applied MFRE-Seq to numerous bacterial and archaeal genomic DNA preparations and successfully confirmed known motifs and identified novel ones. This method should be a useful complement to existing methodologies for studying prokaryotic methylomes and characterizing the contributing methyltransferases.

## INTRODUCTION

DNA can be methylated, that is enzymatically modified with a methyl group, at one of three common positions on the base, converting cytosine to 5-methylcytosine (m5C) or N4-methylcytosine (m4C), or adenine to N6-methyladenine (m6A) [reviewed in (1)]. Methylation is directed by DNA methyltransferases (MTases), each of which catalyzes the formation of one of these three modifications. To a greater or lesser degree, MTases require additional conserved sequence around the methylated base for successful DNA binding and catalysis. These short, conserved sequence elements, often referred to as *motifs* (2), can be deduced by examination of multiple instances of methylation. The distribution of methylation, structure of MTase motifs, and biological functions of DNA methylation differ significantly between prokaryotes and eukaryotes.

In eukaryotes, DNA methylation is associated with control of gene expression, genomic imprinting, X-chromosome inactivation, and RNA splicing (3, 4). While it was long believed that m5C was the only methylated base in eukaryotic DNA, recent studies have confirmed the existence of m6A as well (5–9). Eukaryotic DNA MTases exhibit little intrinsic sequence specificity around the methylated base, acting at weakly specific motifs such as <u>C</u>G in mammals, <u>C</u>G, <u>C</u>HG and <u>C</u>HH in plants (10–12), and V<u>A</u>T in early-branching fungi (8). However, only a subset of bases in these contexts is actually methylated, since eukaryotic MTases are directed to sites of action by other proteins, restricted by the accessibility of chromatin in a given region, or intended to maintain an epigenetic pattern by converting hemi-methylated sites to fully methylated sites following replication. As a result, in eukaryotes sequence context is only one of several factors that determine whether or not a particular base is methylated at any given time.

In bacteria and archaea, all three types of methylation are common. Unlike in eukaryotes, where the bulk of DNA methylation activity is intimately linked with chromosome replication, prokaryotic DNA methylation occurs independently of replication. In prokaryotes, methylation often occurs as part of restriction-modification (R-M) systems, where it protects the chromosome from the action of the cognate restriction endonuclease (REase) (1). However, there are also MTases unaccompanied by REases, so-called orphan or solitary MTases, which can have other biological functions. The most well studied of these are Dam, found in Gammaproteobacteria and involved with mismatch repair, chromosome replication timing, and gene expression (13); Dcm, found in enteric bacteria and involved with gene expression and drug resistance (14); and CcrM, found in Alphaproteobacteria and involved with the regulation of cell cycle and division (15). With the notable exception of some non-specific phage-encoded MTases, microbial MTases tend to have well-defined sequence motifs, typically ranging in length from 4 to 8 bases (16). In microbial genomes, in contrast to those of eukaryotes, most instances of a given motif are methylated, and in fact this fraction often approaches 100%. Nonetheless, it has been observed that a small number of Dam MTase (GATC) sites in *Escherichia coli, Salmonella bongorii*, and *Photorhabdus luminescens* are consistently unmethylated, suggesting competition between MTases and other DNA binding proteins at these few loci (17–20).

Determination of MTase recognition sites was at one time a very tedious process that typically involved installing radiolabeled methyl groups, performing partial digestion of methylated DNA, separating fragments by electrophoresis or chromatography, following the radiolabel, and reconstructing the motif based on analysis of various radiolabeled digest products [e.g., (21)]. As a result, motifs of microbial MTases in R-M systems were often assumed to be the same as those of their cognate REases (for which motifs were significantly easier to determine), as opposed to determined directly. In recent years, however, the direct determination of MTase motifs has become significantly easier with the development of several new technologies.

Single-molecule real-time (SMRT) sequencing from Pacific Biosciences (PacBio) revolutionized the study of m6A and m4C MTases when this technology was introduced in 2010 (22). It was discovered that the polymerase used in this sequencing-by-synthesis method took longer, on average, to incorporate nucleotides opposite these methylated bases than their unmodified counterparts. The presence of m6A and m4C could therefore be inferred by a statistically significant delay in incorporation at a particular locus. The PacBio software environment includes the program MotifMaker (https://github.com/PacificBiosciences/MotifMaker), which extracts a sequence window around each putative methylated base and uses a branch-and-bound search to identify conserved motifs within the set. SMRT sequencing has enabled the determination of motifs for hundreds of new m6A and m4C MTases and has been particularly valuable for studying Type I and Type III R-M systems, for which DNA cleavage patterns cannot be used to deduce binding and methylation motifs (16).

In contrast to m6A and m4C, the SMRT sequencing kinetic signal associated with m5C is significantly weaker, is diffused among several bases around the methylated site, and tends not to be on the methylated base itself (23). Although m5C sites can occasionally be detected with sufficiently high sequence coverage (22, 24, 25) or with hypermodification of the original m5C using TET enzyme (23, 26), on the whole, SMRT sequencing is less suited to the reliable identification of m5C motifs than to m4C or m6A. Other methods of analyzing m5C in DNA have been developed, particularly bisulfite sequencing (27), often considered the “gold standard” of m5C methylation analysis. Treatment of DNA with sodium bisulfite deaminates unmodified cytosine residues to uracil, while leaving m5C residues (and to a lesser extent, m4C residues) intact. Comparison of sequence data from treated and untreated samples enables the distinction of methylated cytosine residues (read as cytosine in both samples) from unmethylated residues (read as thymine in treated samples and cytosine in untreated samples). Whole-genome and targeted bisulfite sequencing have been used successfully to study Dcm modification in *E. coli* (28) and to characterize new m5C methyltransferase motifs in *Enterococcus faecalis* (29), *Enterococcus faecium* (30), and *Prevotella intermedia* (31). In most of these cases, the motif was known or suspected based on other lines of evidence such as SMRT sequencing, but in the case of *E. faecium* the motif was determined *de novo* using MEME (32) motif searching of windows around cytosines protected from bisulfite conversion (30).

While bisulfite sequencing is a powerful technique, it is laborious and can be challenging for those not experienced with it. A balance must be struck between maximizing C-to-U conversion (thereby keeping false positives to a minimum) and minimizing the DNA degradation concomitant with bisulfite treatment. As a result of this damage, relatively large amounts of sample must be used to achieve coverage sufficient to accurately call unconverted C residues. In addition, data analysis can be challenging, requiring specialized alignment tools to map bisulfite-converted reads to the reference sequence.

Some techniques have been developed around interrogating m5C sites using Type IV REases. While “typical” REases cleave unmethylated DNA and are blocked by methylation of the recognition site, Type IV enzymes cleave only when the recognition site bears a specific methyl group and do not cleave at unmethylated sites (2). One particularly useful family of Type IV REases is typified by MspJI (33), and we refer to enzymes of this type as MFREs (<u>M</u>spJI-family <u>RE</u>ases). All MFREs recognize highly degenerate motifs bearing m5C (such as <u>C</u>NNR, the recognition site of MspJI itself), introduce double-strand breaks at a fixed distance 3’ to the m5C base, and leave 4-base 5’ overhangs. The site of cleavage is therefore indicative of the presence of m5C a fixed distance away. In addition, a fully methylated site (that is, a typically palindromic recognition site that is methylated on both strands) will induce double strand breaks on both sides of the site, excising a DNA fragment with the REase motif centered within it and the methylated bases at predictable locations.

MFREs have been used in diverse applications such as random fragmentation (34), measuring changes in methylation by qPCR (35), and quantitation of hm5C using hybridization chain reaction (36). However, the last property above––the excision of fully methylated DNA sites as small DNA fragments with the methylated bases roughly in the middle––immediately suggested the first described application of these enzymes, namely mapping fully methylated sites in eukaryotic genomes at single-base resolution (33, 37, 38). In this work, we have exploited the same property to characterize the recognition sites of microbial m5C MTases, a technique that we term “MFRE-Seq.” Because it relies, not on the identification of any specific site, but merely on the collection of a sufficient number of examples to derive a common sequence signature, it is perhaps an even more straightforward application of MFREs than the mapping of eukaryotic sites. We present the results of analyzing numerous microbial genomes and demonstrate MFRE-Seq as a useful and cost-effective alternative to both bisulfite and SMRT sequencing for characterizing microbial m5C MTases.

## MATERIALS AND METHODS

### Methyl-dependent digestion of DNA

In a typical reaction, 1 µg of genomic DNA was digested in a 40 µl volume of 1x CutSmart buffer with 1 µl of MFRE (MspJI or FspEI; New England Biolabs, Ipswich, MA) and 1.4 µl activator oligonucleotide (15 µM stock; see below) at 37°C for 3 hrs. DNA was subsequently size-selected and purified using either gel or column purification. For gel purification, samples were run on 20% polyacrylamide gels in 0.5x TBE and stained with SYBR Gold. Bands in the 20-40 bp range were excised, and each was placed in a 0.5 ml microcentrifuge tube with a small hole in the bottom. These were placed inside 1.5 ml tubes and centrifuged 5 min at 16,000 x g. DNA was soaked out of the fragmented gel in 100 µl 0.5x TBE buffer 4°C overnight, gel fragments were pelleted by centrifugation, and the supernatant (∼60 µl) retained.

For column purification, smaller DNA fragments (<100 bp) were purified using the Monarch PCR Purification Kit (New England Biolabs) with a modified protocol. Briefly, 2 volumes of binding buffer were added to the digested DNA, and the sample was mixed and applied to the column supplied with the kit. The column was discarded, 2 volumes of 95% ethanol were added to the flow-through, and the sample was again mixed and applied to a fresh column. The column was washed as per the manufacturer’s instructions, and the sample was eluted in 30 µl of 10 mM Tris pH 8.0, 0.1 mM EDTA (TE). DNA was quantitated on a Qubit fluorimeter (Life Technologies, Eugene, OR), and a typical yield was < 15 ng.

### Preparation of enzyme activator oligonucleotides

A standard 28 base methylated hairpin oligonucleotide (CTGC<u>C</u>AGGATCTTTTTTGATC<u>C</u>TGGCAG) is provided by the manufacturer (New England Biolabs) at 15 µM. We also designed three modified derivatives of this oligonucleotide (Integrated DNA Technologies, Coralville, IA; see Results). Activator-U and activator-NU replaced the 6-base poly-dT run at the hairpin loop with a 6-base poly-dU run. Activator-N and activator-NU attached an amino modifier C6 at the 5’ end of the oligo (i.e., 1-aminohexane attached via C6 to the 5’ phosphate) as aligation blocking group. These modified derivatives were resuspended to 15 µM, denatured at 95° for 5 min and cooled slowly to room temperature to anneal the hairpins.

### Library preparation and sequencing

Libraries were prepared using the NEBNext Fast DNA Library Prep Set for Ion Torrent (New England Biolabs) with IonXpress Barcode Adapters (Ion Torrent, Carlsbad, CA), or the NEBNext Ultra II DNA Library Prep Kit for Illumina and NEBNext Multiplex Oligos for Illumina (both New England Biolabs).

Ion Torrent libraries were prepared using 18 µl (∼4 ng) gel-purified DNA or 25 ng column-purified DNA as input according to the manufacturer’s protocol, with the following exceptions. To minimize the denaturation of the small DNA fragments that would occur with a heat-treatment step, end repair was carried out with a mixture of bead-immobilized T4 DNA polymerase and T4 polynucleotide kinase (both kind gifts of Dr. M. Xu, New England Biolabs) in a 20 µl volume of End Repair buffer, 20 min at 25°C. The immobilized enzymes were removed on a magnetic rack.Barcode and P1 adapters were used at 1:100 dilutions. Following adapter ligation, the library was amplified with 10-12 cycles of PCR.

Illumina libraries were prepared using 25 ng column-purified DNA as input according to the manufacturer’s protocol. Adapters were used at 1:10 dilutions, and the library was amplified with 5 cycles of PCR. Fragment sizes were determined using a BioAnalyzer (Agilent Technologies, Santa Clara, CA), and libraries with fragments over 500 bp were size-selected with the following protocol. 50 µl of library was mixed with 0.55x NEBNext Sample Purification Beads (New England Biolabs; “beads”), incubated at room temperature for 5 min, and the supernatant was retained. The supernatant was mixed with 0.9x beads and incubated at room temperature for 5 min; beads were washed twice with 200 µl 80% ethanol, dried 5 min, and DNA fragments were eluted in 30 µl TE. Multiplexed samples were sequenced on a MiSeq (Illumina, San Diego, CA) using a 2×50 paired-end kit.

### Sequence read processing

Paired-end Illumina reads were merged using SeqPrep (https://github.com/jstjohn/SeqPrep) with a minimum overlap for merging of 20, a minimum overlap of 5 for adapter trimming, and a minimum 26 bp merged read size. Ion Torrent reads were merged and trimmed by the manufacturer’s software. A set of Perl scripts performed the subsequent mapping, motif-finding, and motif analysis functions. Trimmed reads were mapped to a reference sequence, and only exact matches over the entire read length were retained. Duplicate reads were collapsed, and the non-redundant set of reference-matching reads was binned by length. Motif-finding was performed on each bin separately, typically in the range of 26-45 bp.

### Relationship between methylation and read length

MFREs cleave at a fixed distance (16 bp 3’) from the m5C base and require m5C residues on both strands to generate fragments of defined length. Consequently, the precise fragment length generated by cleavage of a fully methylated motif depends on the relative positions of the m5C on the two strands (Figure 1). If *x* is the number of bases that separate the top-strand m5C in the motif from the bottom-strand m5C (for examples, *x* = 1 for A<u>GC</u>T and *x* = –2 for C<u>C</u>W<u>G</u>G), then the resulting fragment length from MFRE cleavage will be *l*_*x*_ = *x* + 33 provided the DNA was cut at the expected distance on both sides. Sequence reads derived from these fragments will also be of length *l*_*x*_ provided the sequence was also accurately trimmed *in silico* by the adapter-trimming algorithm. For an 8-base motif (the longest observed m5C-containing motif), *l*_*x*_ can vary from 26 to 40 bp, and so we typically examine reads in this length range. The value of *x* cannot be 0 (so *l*_*x*_ cannot be 33), since this would mean the top and bottom strand m5C residues would be base-paired with each other.

**Figure 1.**
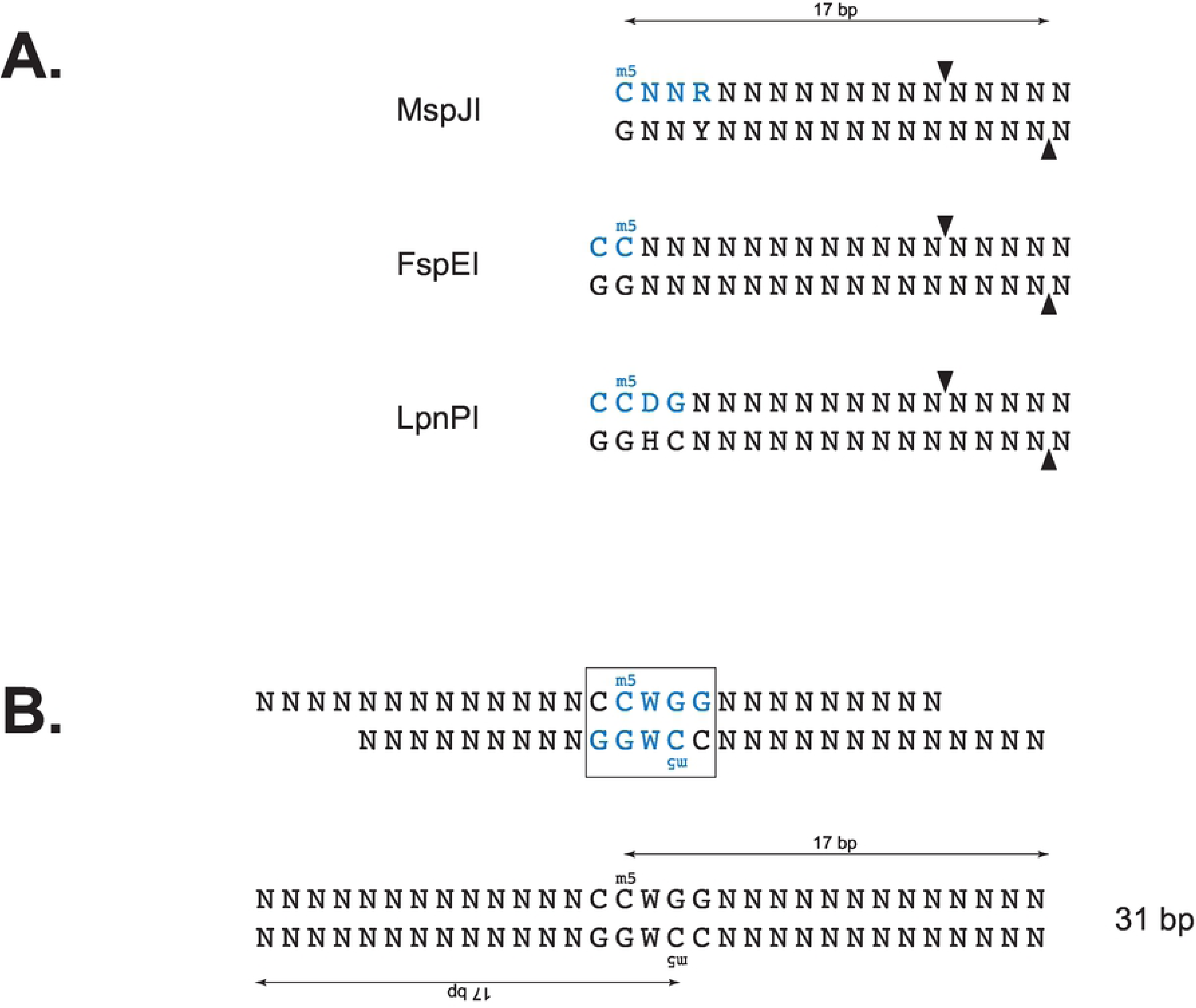
MFRE cleavage and formation of library inserts.A) Recognition sites (blue) and cleavage positions (arrows) of three commercially available MFREs. B) Product of MspJI (recognition site in blue) cleavage of the fully methylated motif C<u>C</u>W<u>G</u>G (boxed), before and after end repair. The m5C residue on each strand is the 17^th^ base from the 3’ end.

In order to determine what fraction of reads are correctly cut and trimmed, we analyzed samples where the motif and specific methylated bases were known *a priori*. In such cases, it is helpful to represent a motif-containing read as a pair of lengths, (*d*_1_, *d*_2_), where *d*_1_ and *d*_2_ are the “cleavage distances” between the methylated bases and the ends of the read. Strand orientation is random, so by convention we represent the pair such that *d*_1_ ≤ *d*_2_. These lengths are measured from the top-strand m5C to the 3’ end of the read and from the G opposite the bottom-strand m5C to the 5’ end of the read. For a correctly cut and trimmed read, *d*_1_ = *d*_2_ = 16, and such a read would be designated “(16,16)”. We typically group values of *d* less than 14 and greater than 18 as “–13” and “19+”, respectively.

### Enriching for methylation-derived reads

Sequence reads that are derived from MFRE cleavage at the standard distance (16 bp 3’ to the m5C on both strands) and accurately adapter-trimmed *in silico* we term “CCMD reads” (<u>c</u>orrectly <u>c</u>ut and trimmed <u>m</u>ethyl-<u>d</u>erived reads). It is from CCMD reads that we can easily determine Mtase motifs by direct alignment. However, the sequence reads produced by MFRE-Seq contain many types of reads besides CCMD reads. These can include sequences derived from contaminants; error-containing sequences derived from the intended DNA; sequences from inserts broken on one or both sides by mechanical or non-MFRE enzymatic breakage; interstitial fragments between methylated fragments; and MFRE-cleaved fragments that were not correctly cut and trimmed. The last of these categories in particular is expected to be numerous, since MFREs have been shown to occasionally cut 1 bp beyond the expected length. We therefore included several *in silico* filtering steps to enrich the data set for CCMD reads.

By definition, all CCMD reads are (16,16), meaning they have a C as the 17^th^ base from the 3’ end and a G as the 17^th^ base from the 5’ end. The converse is not always true, since some reads (1 in 16 in a random set) will coincidentally have a C and a G at these respective positions. However, by filtering out all reads that do not satisfy these properties, we significantly enrich for CCMD reads in our sequencing data. This process is effective for many but not all motif and methylation structures (Table S1). Thus, there are three filtering steps altogether: (1) removal of reads that are not exact matches to the reference (“reference-filtering”), (2) removal of reads outside the 26-40 bp range (“size-filtering”), and (3) removal of reads not containing C and G at the expected positions (“base-filtering”).

### Deduction of motifs from read sets

To look for motifs, filtered reads of a given length were aligned in an ungapped fashion, and for each column of the alignment the nucleotide distribution was determined. Using Kullback-Leibler (KL)-divergence, this distribution was compared to the distributions expected of all possible IUPAC base symbols (degenerate and non-degenerate) based on the actual overall base frequencies of the reference sequence. The IUPAC symbol with the smallest KL value was assigned to that column. Consecutive N’s at the start and end of the alignment were removed, and the string of remaining symbols was scored for complexity. All strings have at least one C and one G due to the base-filtering criterion, so the string had to have sufficient additional complexity to be considered a possible motif. Individual IUPAC symbols were assigned the following arbitrary scores, and the score for the string was the sum of all individual base scores: (A, C, G, T) = 8; (R, Y, M, K, S, W) = 4; (H, B, V, D) = 2. One C, one G, and all N’s were removed from the string prior to scoring, and all strings with complexity scores ≥ 10 were considered candidate motifs. Thus, CCGG (score = 16) would be considered a candidate motif, but CRYG (score = 8) and CNNG (score = 0) would not.

### Deconvolution of composite motifs

The approach above oversimplifies certain cases, requiring additional analysis. These include read sets with multiple motifs, read sets with non-palindromic motifs, and motifs with base dependencies. Such cases often present themselves as candidate motifs with degenerate bases, yet with enough complexity to pass the scoring threshold above. Degenerate bases in a motif can result from legitimate tolerance for multiple bases at a given position in the MTase’s DNA binding footprint, or they can result from the inappropriate merging of independent substrate sequences.

| Type | True Motif(s) | Apparent Motif |
| --- | --- | --- |
| Multiple motifs | <u>C</u>CGG, <u>C</u>AT<u>G</u> | <u>C</u>M<u>K</u><u>G</u> |
| Non-palindromic | C<u>C</u>C<u>G</u>C / G<u>C</u>G<u>G</u>G | S<u>C</u>S<u>G</u>S |
| Base dependency | G<u>C</u>TAGC + G<u>C</u>C<u>G</u>G<u>C</u> | G<u>C</u>YR<u>G</u>C |
| True degeneracy | G<u>C</u>YR<u>G</u>C | G<u>C</u>YR<u>G</u>C |

To distinguish these possibilities, degenerate motifs are further analyzed by examining the frequencies of all non-degenerate instances of that motif among filtered reads of the appropriate length. Cases of inappropriate merging will become apparent as certain base combinations within the degeneracy do not appear or appear very rarely. For the two apparent GCYRGC cases above, in the top case only TA and CG would appear at appreciable frequencies at the YR positions, while in the bottom case all four combinations (TA, CG, TG, and CA) would appear at similar frequencies.

## RESULTS

### Experimental design

To determine m5C MTase motifs, we employed the following general approach (Figure 2). Purified genomic DNA was digested with one or more MFREs, and small (<100 bp) fragments were selectively purified using either gel electrophoresis/excision or spin-column binding/elution.

**Figure 2.**
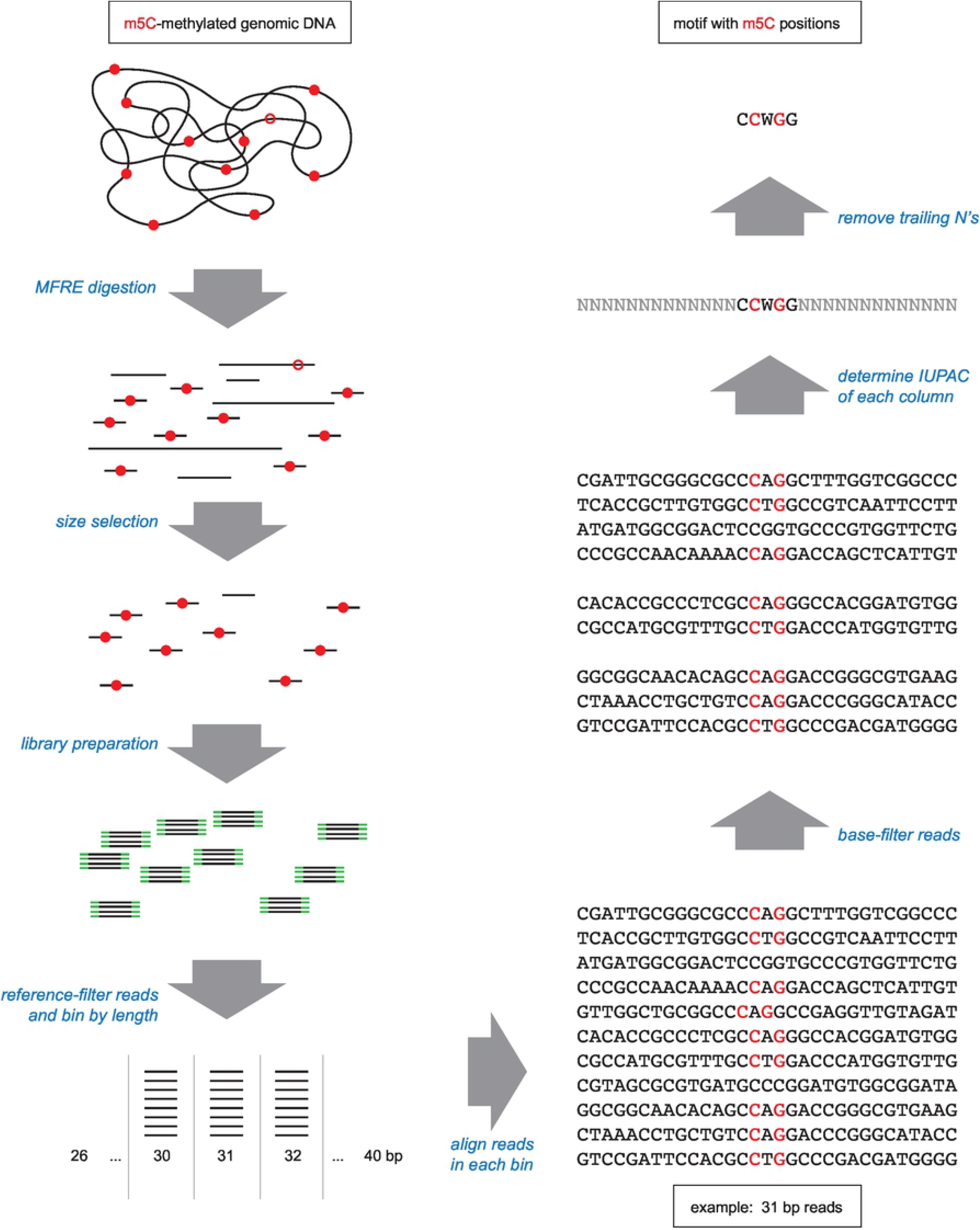
Overview of MFRE-Seq. Genomic DNA containing motifs that are fully methylated (red dots) or hemi-methylated (open red circles) is digested with one or more MFREs. Size selection enriches for the small fragments that result from MFRE cleavage of fully methylated sites, and sequencing libraries are prepared from these fragments (adapters in green). Sequence reads are then mined for motifs. The computational method for doing so described in this work involves binning reads by length, enriching for CCRM reads by base-filtering, aligning, and examining the base distribution at each position.

Sequencing libraries were then constructed from the purified fragments and sequence data obtained. After paired-read merging and adapter trimming, the typical range of read lengths was 20-80 bases for Ion Torrent and 26-80 bases for Illumina. Reads were mapped to a reference sequence, those that were not exact matches to the reference (i.e., no gaps and no mismatches) were discarded, and those that remained were oriented to the top strand of the reference. Remaining reads were then base-filtered (see Materials and Methods) and sorted by length, and the set of reads of each length within the range 26-40 bp was tested separately for the presence of conserved motifs.

### Sequence read analysis from cleavage of the *E. coli* Dcm site

We first tested this approach on several samples where the methylated motif was known by other means, starting with *E. coli* K-12, whose genome is methylated by Dcm at C<u>C</u>W<u>G</u>G sites but is free of other m5C MTases. This site can be effectively cut by both MspJI (as it overlaps <u>C</u>NNR) and FspEI (as it overlaps C<u>C</u>), and the expected (16,16) cleavage products are 31 bp. We digested 1 µg of genomic DNA from *E. coli* DHB4 (an F^+^ K-12 derivative) with MspJI or FspEI and prepared Illumina sequencing libraries from 25 ng of column-purified digest, which was subsequently size selected (see Materials and Methods for details) and sequenced with 2×50 paired-end kits. Four duplicate libraries constructed from separate digests were sequenced, one multiplexed with a second sample and run on a MiSeq and the other three each multiplexed with eight other samples and run on a NextSeq. In all four trials, the fraction of all reads that were exact matches to the reference sequence was >96% (Table 1). The mean copy number of each read varied between trials, from 15x to 80x (Table 1).

**Table 1.** Replicate experiment statistics, *E. coli* DHB4 genomic DNA.

| Replicate | 1 | 2 | 3 | 4 |
| --- | --- | --- | --- | --- |
| MFRE | FspEI | FspEI | FspEI | MspJI |
| Platform | MiSeq | NextSeq | NextSeq | NextSeq |
| Multiplex | 2 | 9 | 9 | 9 |
| Pairs Merged<sup>a</sup> | 7,925,782 | 15,288,614 | 19,851,338 | 13,396,381 |
| Pairs with Adapters<sup>a</sup> | 7,118,719 | 14,941,064 | 19,027,975 | 13,258,050 |
| Pairs Discarded<sup>a</sup> | 270,585 | 312,512 | 600,062 | 550,688 |
| Reads Matching Reference<sup>b</sup> | 7,657,629 | 14,944,470 | 19,419,069 | 13,099,298 |
| Fraction Matching Reference | 0.966 | 0.977 | 0.978 | 0.978 |
| Unique Reads Matching Ref. | 191,145 | 230,260 | 243,416 | 876,198 |
| Mean Redundancy | 40 | 65 | 80 | 15 |
| Unrepresented CCWGG sites | 696 | 684 | 663 | 636 |

<sup>a</sup> Output from SeqPrep.
<sup>b</sup> Exact matches, no polymorphisms or indels.

Knowing the motif and methylated bases ahead of time, we could classify the sequence reads containing CCWGG in terms of the distances between the methylated bases and the ends of the read. (If a read contained multiple instances of the motif, we chose the motif instance closest to the center of the read for this purpose.) Tables 2a and 2b show the number of all reads and base-filtered reads, respectively, with each possible pair of distances. Among all (reference-matching) reads, the most common categories were (16,16) > (16,17) >> (16,19+) > (17,17) > (15,16) >> all other categories (Table 2a). Thus, the vast majority of reads were either (16,16) reads (length 31; 67.6%) or (16,17) reads (length 32; 23.4%). Combined, 96.4% of reads have at least one flank that was correctly cut and trimmed (*d* = 16), but a sizeable number (26.5%) had at least one flank with 1 extra base. The extra bases likely result from the MFREs cutting 1 base farther from the m5C than expected, as has been observed previously (39).

**Table 2a.**
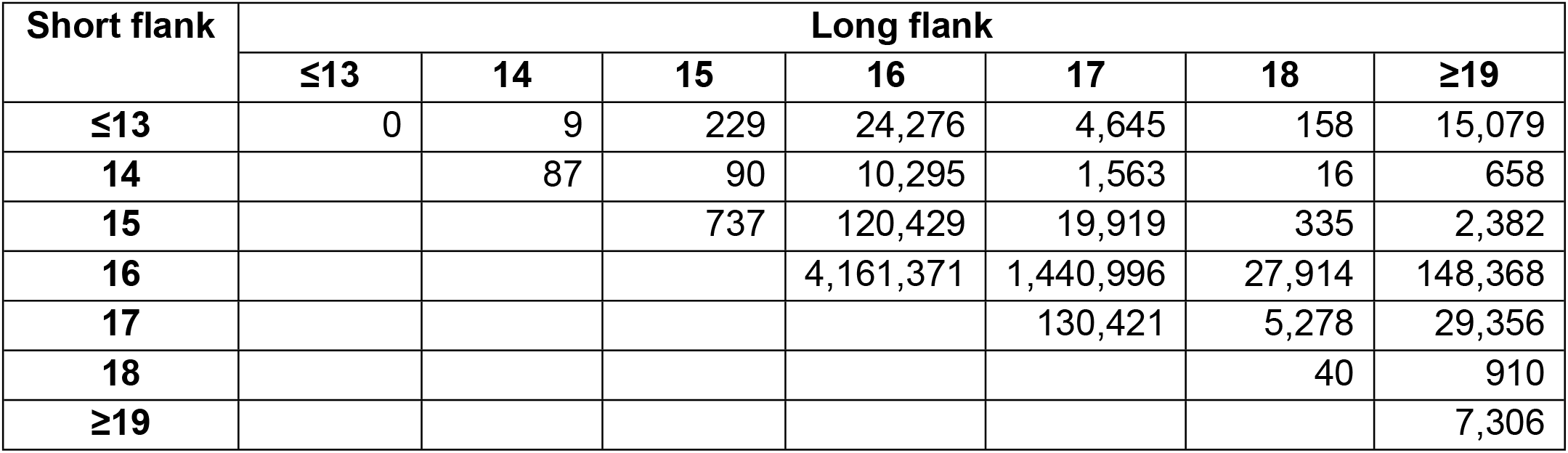
Flank length analysis of all reference-matching, motif-containing reads derived from Illumina DHB4 run.

| Short flank | Long flank |  |  |  |  |  |  |
| --- | --- | --- | --- | --- | --- | --- | --- |
|  | ≤13 | 14 | 15 | 16 | 17 | 18 | ≥19 |
| ≤13 | 0 | 9 | 229 | 24,276 | 4,645 | 158 | 15,079 |
| 14 |  | 87 | 90 | 10,295 | 1,563 | 16 | 658 |
| 15 |  |  | 737 | 120,429 | 19,919 | 335 | 2,382 |
| 16 |  |  |  | 4,161,371 | 1,440,996 | 27,914 | 148,368 |
| 17 |  |  |  |  | 130,421 | 5,278 | 29,356 |
| 18 |  |  |  |  |  | 40 | 910 |
| ≥19 |  |  |  |  |  |  | 7,306 |

**Table 2b.** Flank length analysis of base-filtered reference-matching, motif-containing reads derived from Illumina DHB4 run.

| Short flank | Long flank |  |  |  |  |  |  |
| --- | --- | --- | --- | --- | --- | --- | --- |
|  | ≤13 | 14 | 15 | 16 | 17 | 18 | ≥19 |
| ≤13 | 0 | 0 | 55 | 7,948 | 0 | 0 | 1,466 |
| 14 |  | 0 | 15 | 1,160 | 0 | 0 | 2 |
| 15 |  |  | 737 | 120,429 | 0 | 0 | 425 |
| 16 |  |  |  | 4,161,371 | 0 | 0 | 38,066 |
| 17 |  |  |  |  | 0 | 0 | 0 |
| 18 |  |  |  |  |  | 0 | 0 |
| ≥19 |  |  |  |  |  |  | 372 |

Because not all reads were of the categories above, and not all contained the CCWGG motif, we compared the reads we obtained (from the trial in column 1 of Table 1) to a theoretical FspEI digest of *E. coli* DHB4. We classified the theoretical fragments as one of six categories: “motif-cleaved” (when exactly cut, these are CCMD fragments), “interstitial” (regions between motif-cleaved fragments), “overlap-short” (non-motif containing fragments created by cutting two CCWGG sites spaced less than 30 bp apart), “overlap-long” (containing 2 motifs, created by cutting CCWGG sites less than 30 bp apart without cutting between them), “other” (created by more complicated situations such as 3 or more clustered motifs), and “concatenated” (reads spanning an expected cut site, which most often consist of a motif-containing CCMD fragment joined to an interstitial fragment). The real reads were matched to the theoretical fragments and classified as one of four categories based on how they were cut: “exact” (cutting at the expected MFRE site on both ends), “approximate” (cutting within 4 bp of the expected site on both ends), “one-cut” (cutting within 4 bp of the expected site on one end only), and “neither” (not cutting within 4 bp of the expected site on either end). Results are shown in Table 3.

**Table 3.** Comparison of real sequence reads with theoretical digest fragments of *E. coli* DHB4.^a^

|  | **Exact** | **Approximate** | **One-Cut** | **Neither** |
| --- | --- | --- | --- | --- |
| **Motif-Derived** | 11,731<br>*8,940,794*<br>(762x) | 64,631<br>*3,844,900*<br>(59x) | 1,651<br>*10,802*<br>(6.5x) | 0<br>*0*<br>(n/a) |
| **Interstitial** | 0<br>*0*<br>(n/a) | 5,718<br>*1,205,541*<br>(211x) | 76,774<br>*329,449*<br>(4.3x) | 20,182<br>*31,260*<br>(1.5x) |
| **Overlap-Short** | 24<br>*3,429*<br>(143x) | 3<br>*104*<br>(35x) | 0<br>*0*<br>(n/a) | 0<br>*0*<br>(n/a) |
| **Overlap-Long** | 291<br>*6,516*<br>(22x) | 723<br>*6,538*<br>(9.0x) | 1,298<br>*28,434*<br>(22x) | 10<br>*30*<br>(3.0x) |
| **Other** | 0<br>*0*<br>(n/a) | 6,938<br>*350,915*<br>(51x) | 1,721<br>*9,972*<br>(5.8x) | 101<br>*168*<br>(1.7x) |
| **Concatenated** | 0<br>*0*<br>(n/a) | 0<br>*0*<br>(n/a) | 18,891<br>*141,864*<br>(7.5x) | 19,573<br>*33,754*<br>(1.7x) |

<sup>a</sup> For each category, the top line (Roman type) shows the number of unique sequence reads, the middle line (italic) shows the number of all sequence reads, and the bottom line (in parentheses) shows the copy number of the reads in this category (all/unique).

The relatively low numbers of concatenated fragments, as well as the low number of motif-containing fragments cut on only one side, indicate that the digest was largely complete (Table 3).The vast majority of reads were motif-cleaved, either exactly or approximately cut on both sides. The lower copy number of approximately cut sequences here reflects the fact that approximate cutting generates a variety of ends on both sides. When CCWGG cutting sites overlapped (within 30 bp of each other), the most common fragments were of the overlap-long type, perhaps indicating that MFREs have a harder time cutting shorter fragments. In other words, once an overlap-long fragment is created, the enzyme has a harder time cutting it further. Most theoretical interstitial fragments are longer than 80 bp, so many of those that appear in the sequence reads are cut on one or neither side by an MFRE and must be broken on the other side by some other process. Those less than 80 bp are present at high copy number, but all of these are approximately rather than exactly cut.

In addition to reads without the motif, we also observed CCWGG sites in the reference without a corresponding read in the sequence data. In our four replicate *E. coli* DHB4 experiments, we observed a mean of 670 out of 12,321 (5.4%) CCWGG sites to be unrepresented by either (16,16) or (16,17) reads in our sequence data (Table 1). While some of these may be truly unmethylated, alternative explanations for the non-representation of these sites include errors in the reference sequence; lack of full methylation at specific sites due, for example, to steric hinderance by DNA binding proteins; systematic cleavage bias away from (16,16) products; and interference of closely proximal sites.

We compared the four sets of unrepresented sites and found that the vast majority of them (623) were common to all four data sets (Figure S1). There was no significant distinction between either the MFRE used for the digest or the machine used for sequencing. We mapped the locations of these 623 sites and found they corresponded to repeat regions, most notably a region of the chromosome duplicated on the F’ element and the rRNA gene clusters. The apparent absence of these sites is therefore due to the pileup of reads derived from duplicate locations on the chromosome to a single locus. After filtering out these repeat locations, we found only 100 of 12,321 sites (0.8%) of CCWGG sites unrepresented in the data.

### Motif finding approach

The results in Table 2a show that many reads are not of the (16,16) variety, and so the precise motif location in any given read cannot be assumed with certainty. Table 2b shows that, in this particular instance at least, base filtering effectively enriches for (16,16) reads: there are 4,161,371 reads of type (16,16), representing 67.6% of all reads and 96.0% of base-filtered reads.

We looked for data features independent of the knowledge of motif content or structure that would be useful for inferring whether or not reads of a given length are CCMD reads. In particular, we looked at the number of sequencing reads obtained, the redundancy (copy number) of reads, and the fraction of reads that survived base filtering. All reads, even those generated by random processes, may potentially appear multiple times in the data due to amplification during library preparation. However, because CCMD reads are generated by repeated cleavage at a limited number of genomic locations, we would expect all three of these metrics to be significantly higher for CCMD reads than for the “background” consisting of reads generated by more random processes. Since we expect CCMD reads to be in the 26-40 base range, we use as background values of these metrics the mean values calculated for reads outside of this range (46-80 bases in length).

Figure 3 shows that, for the *E. coli* K-12 DHB4 experiment, the three metrics mentioned above peak sharply around length 31 (the expected (16,16) length). The background rate of redundancy is roughly 13 copies per sequence, while the redundancy at lengths 31 and 32 are significantly higher, at 245 and 67 copies per sequence, respectively (Figure 3A). These two lengths are also those with the highest absolute numbers of reads (Figure 3B). The “background” fraction of reads that survived base filtering was 8.1% (comparable to 6.2%, that expected of randomly generated reads of 50% G+C), while this fraction was significantly higher among reads of length 30 (92%) and 31 (99%) (Figure 3C).

**Figure 3.**
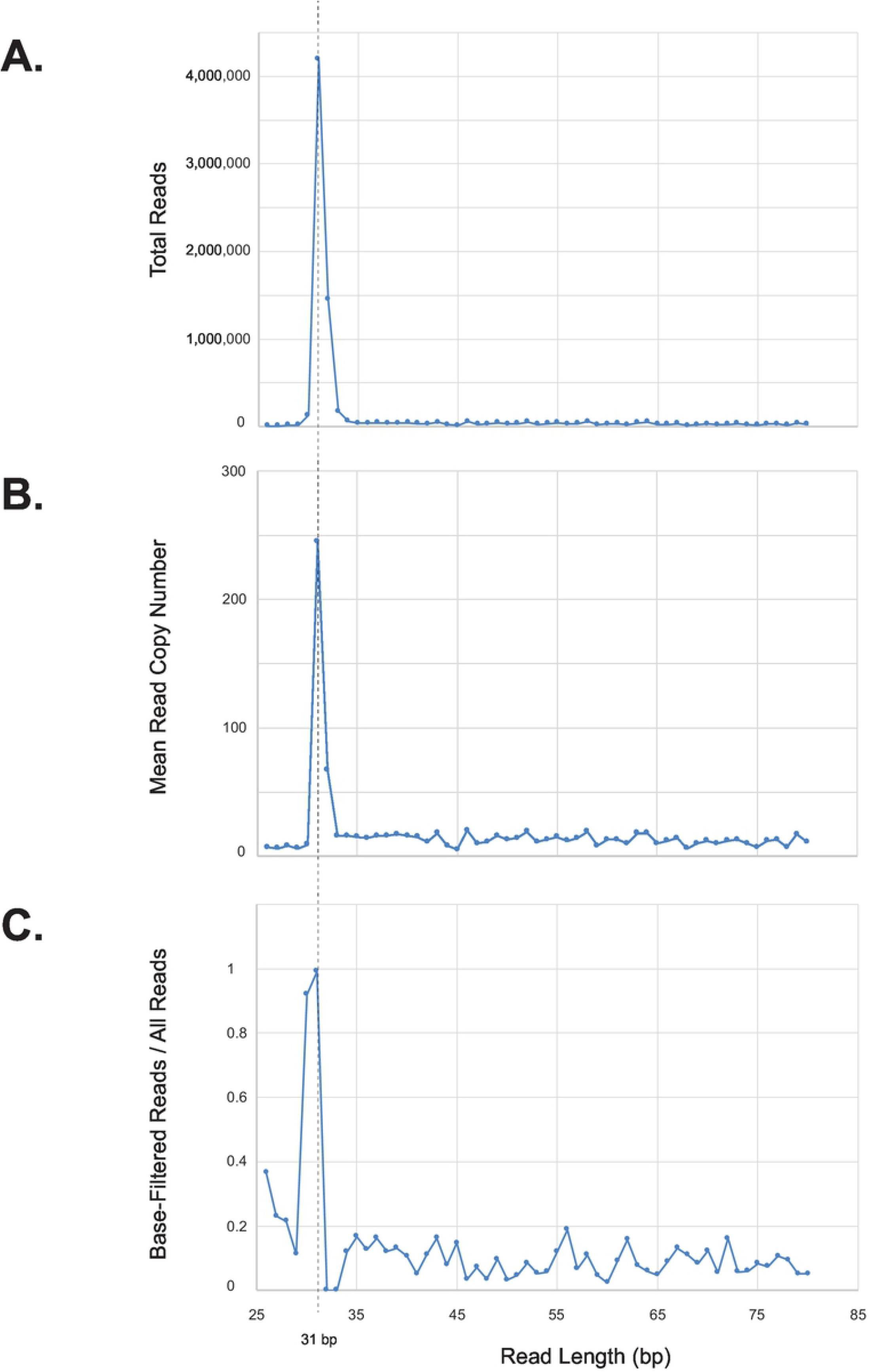
Diagnostic statistics from Illumina sequencing of FspEI-digested *E. coli* K-12 DHB4. This strain is methylated by Dcm at C<u>C</u>W<u>G</u>G sites, resulting in 31 nt CCMD reads (dotted vertical line). All numbers are for reference-matched reads. Top: total number of reads of each length. Middle: mean read copy number of each length. Bottom: fraction of reads of each length that passed base filtering.

The vast majority of reads of length 30 (high number of reads and fraction of base-filtered reads, but low redundancy), 31 (high number of reads, fraction of base-filtered reads, and redundancy), and 32 (high number of reads and redundancy, but low fraction of base-filtered reads) are (15,16), (16,16), and (16,17), respectively (Table S2). We further sorted the reads of length 31 by copy number, and for each copy number we examined the fraction of base-filtered reads and the fraction of (16,16) (i.e., motif-containing) reads. The copy number ranged from 1 to 7129 copies, with a mean of 245. Most of the non-(16,16) reads of this length are (15,17) and are present in less than 15 copies, which is approximately the “background” rate.

The data above suggests the following approach to determining motifs (of unknown sequence, length, and number) in sequencing data from MFRE cleavage:

1. Establish background levels of read numbers, redundancy, and base-filtered read fraction based on reads outside the expected range of true cleavage products (e.g., those from 46-80 bp).
2. Identify read lengths for which at least one of the metrics above (numbers of reads, mean copy number, or fraction of base-filtered reads) is above the background levels.
3. For each of these lengths, eliminate reads with copy numbers at or below the background level, and determine motifs from the remainder. Keep those motifs that are sufficiently specific to be real. Some of these may be spurious, derived from overlapping instances of the same motif in (16,17) or (15,16) reads, and we expect such spurious motifs to be less specific than the true motifs from which they are derived.
4. From the set of possible motifs, identify that with the highest per-base specificity score (*m*_max_) and save it in the final set of motifs. From the length of read from which it is derived (i.e., the presumed (16,16) length), identify the methylated bases.
5. Delete all reads derived from motif *m*_max_ with exact cleavage on at least one end. (This includes not just (16,16) reads but also (*x*,16) and (16,*x*), where *x* ≠ 16).
6. Repeat steps 3-5 on the reduced set of reads, identifying the next most specific motif. Iterate until no more motifs are found.

Applying this pipeline to the *E. coli* DHB4 data set, we obtained a single motif, the expected C<u>C</u>W<u>G</u>G.We have used this same pipeline to determine the other motifs presented in this work.

### Reduction of unproductive sources of sequence

MFRE enzymes require interaction with multiple instances of their recognition sites for efficient cleavage, and so cleavage of a desired substrate can be driven towards completion by the addition of an “enzyme activator” oligonucleotide containing the MFRE’s methylated recognition site (39). The activator provides an excess of recognition sites for binding *in trans* but is too short to be cleaved by the enzyme.

In our initial experiments, sequence data included some reads derived from the MFRE enzyme activator. To prevent sequencing of the activator, we tested three derivatives: “activator-U” (where two adjacent thymine residues in the loop were replaced by uracils, preventing amplification of library molecules derived from it), “activator-N” (where the 5’ phosphate is blocked by a C6-amino modification, preventing ligation with the library adapters), and “activator-UN” (containing both modifications). All three adapters stimulated cleavage of *Pseudomonas mendocina* genomic DNA (G<u>G</u>W<u>C</u>C modified, see Table 4) by MspJI to a comparable degree (Figure S2). We then prepared libraries of *P. mendocina* and *Bacillus* sp. N3536 genomic DNA digested with MspJI with the different activators and sequenced on the Ion Torrent platform. The numbers of reads matching activator-N and activator-NU were close to the background where no activator was added to the reaction. Activator-U provided at least 10x reduction in the number of reads compared to the standard, unmodified activator but more than activator-N and activator-NU (Table S3). We therefore used activator-N in all subsequent library preparation reactions.

**Table 4.**
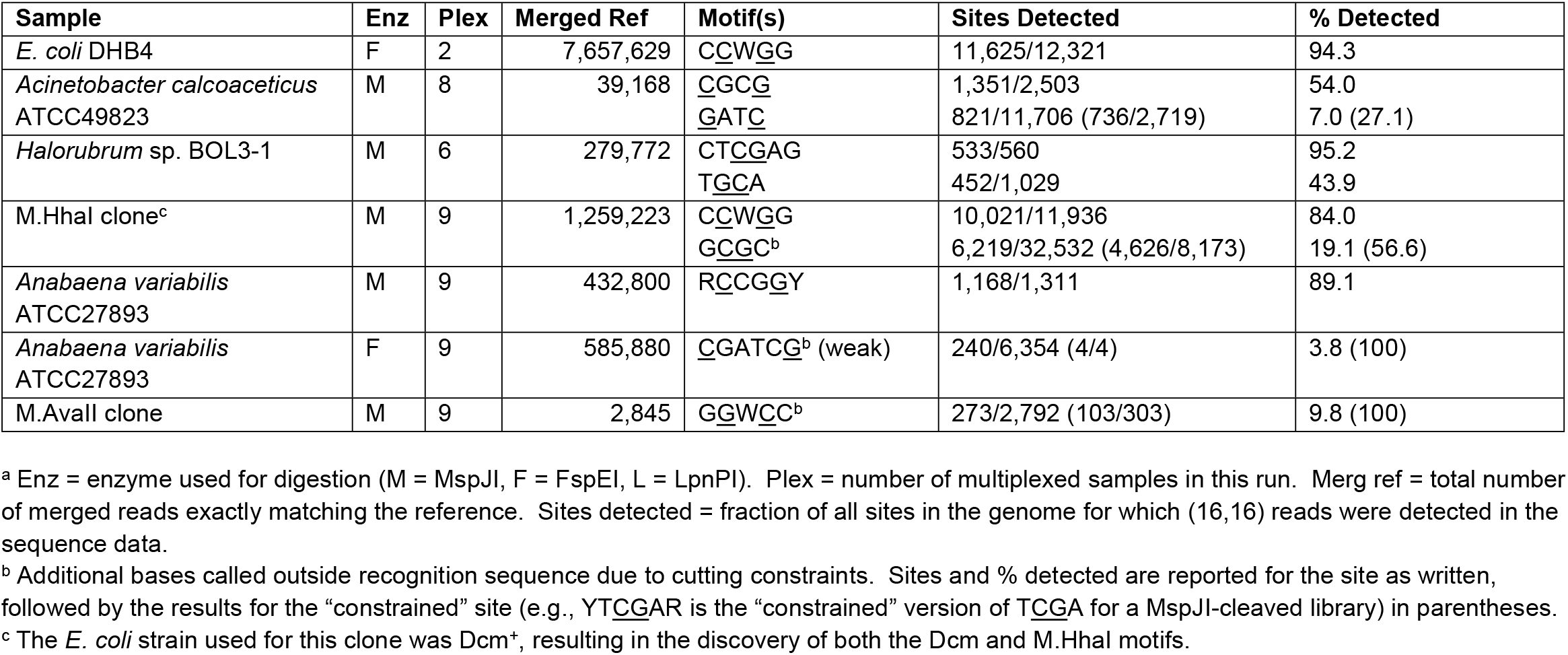
Motifs determined using the Illumina MiSeq platform.^a^

We also tested the distribution of reads obtained using three methods of post-digest fragment purification: gel-purification, a standard column-purification protocol used for oligonucleotide cleanup, and a two-step column-purification protocol more suited to separating oligonucleotides from larger DNA fragments. We examined the distribution of read lengths from three independent experiments, all sequenced on the Ion Torrent platform (Figure S3). Unsurprisingly, the background of non-MFRE-derived reads was significantly higher with the single-column method than the other two methods (Figure S3). Of the other two methods, the two-column purification method is significantly less labor-intensive, requires less time, and does not suffer from contamination with DNA marker-derived reads, we have primarily used this method for digest cleanup prior to library preparation.

### Minimal examples to derive motif

In the above example, there are 12,321 instances of CCWGG in the *E. coli* DHB4 genome, of which 11,940 (96.9%) were represented in the sequence data by (16,16) reads. There may be other experiments in which the motif is comparatively rare, and/or in which fewer reads are generated, so we wished to determine how many examples [i.e., unique (16,16) reads] were required to accurately determine a motif.

We generated randomized sequences *in silico* that included “*in silico* CCMD” sequences [mock (16,16) reads with the motif in the center and flanks of random bases] and a specified fraction of “*in silico* non-CCMD” sequences (identical in length to the “true” sequences, but with randomized sequence replacing all of the motif except the C and G bases required to pass base filtering). All randomized sequence was biased to a predetermined %G+C content, and degenerate positions within the motif were randomly assigned among the permitted bases.

We generated sets of random reads to simulate determination of the C<u>C</u>W<u>G</u>G motif from 31 base reads using the KL-divergence based motif finding script. Adding one read at a time, progressively larger read sets were generated in order to determine the largest number of unique reads from which the program *incorrectly* deduced the motif (if sets of size *a*+1 through *a*+50 all correctly deduced the motif but *a* did not, the experiment stopped and *a* was considered the largest incorrect set). Several experiments were run, varying %G+C between 30-70 and the fraction of non-CCMD reads between 0-0.2. The results of each experiment were determined as the mean of 25 replicate runs.

Results are shown in Figure 4. The method is robust to changes in %G+C and to fractions of non-CCMD reads up to 0.1. With increasing non-CCMD fraction between 0.1-0.2, the number of reads required to correctly deduce the motif rises rapidly, and above 0.2 it was impossible to accurately determine the motif even up to 100,000 unique reads (far above the number typically possible for a bacterial genome). In the *E. coli* DHB4 experiment above, there were 11,064 unique base-filtered reads above the background redundancy level of 13, and only 10 of these were non-CCMD reads, so the non-CCMD fraction in this particular example was 0.001. For 0% non-CCMD reads, correct deduction required 33-69 examples; for 5%, 34-66 examples; and for 10%, 45-102 examples. We obtained similar results for a variety of motifs of varying motif lengths, flank lengths, and degenerate base content (data not shown). Typically, higher numbers of reads were required at more skewed %G+C values due to the lower complexity of these randomized sequences.

**Figure 4.**
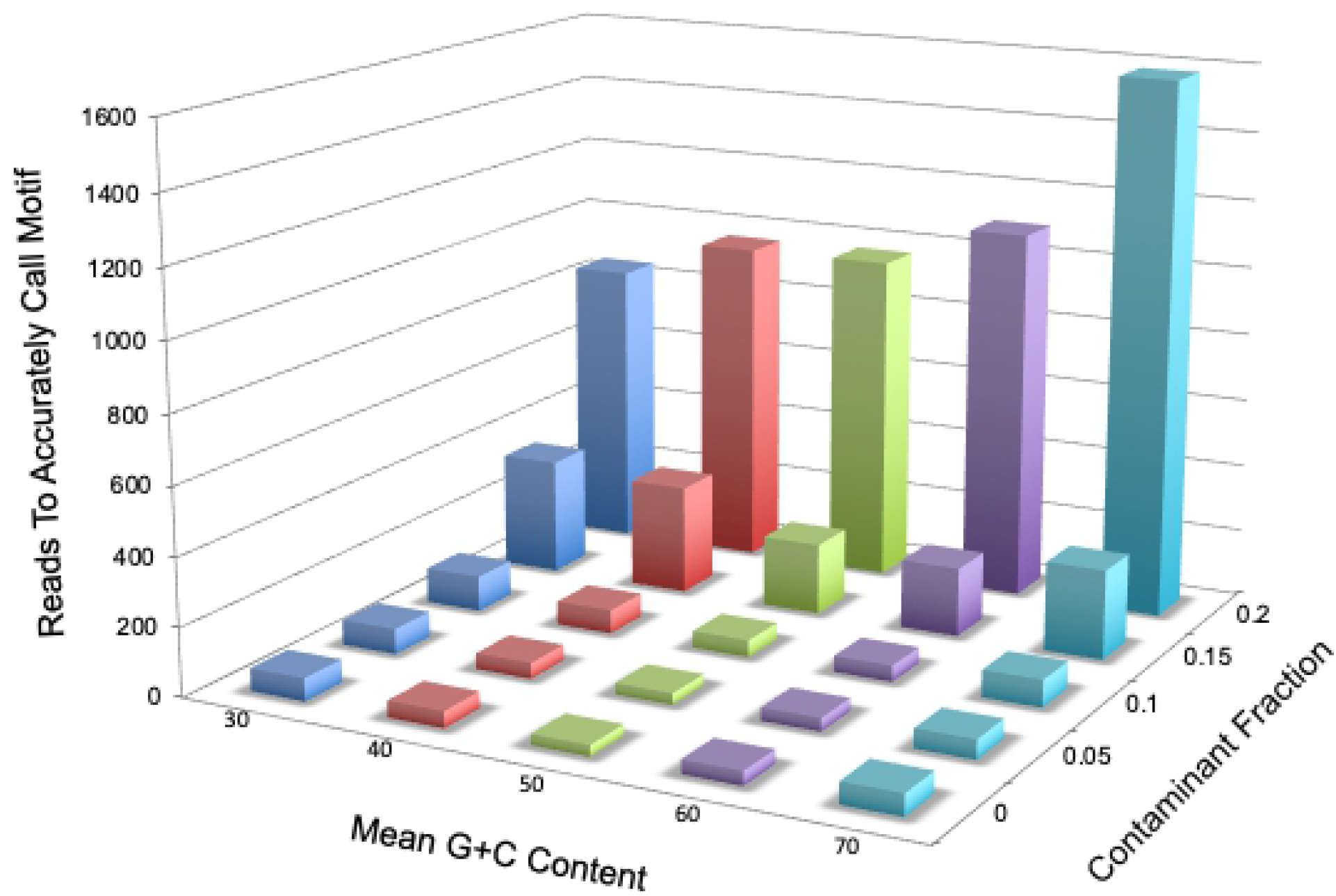
Bar graph of random read analysis. For each combination of G+C content and fraction of non-CCMD reads (horizontal axes), we determined the largest number of reads at which the motif was inaccurately called and added one to this value. The number of reads to accurately call the motif (vertical axis) was calculated as the mean of 25 replicate determinations.

### Characterization of m5C motifs in multiple genomes

We then applied this motif-finding approach to other genomes, using two DNA sequencing platforms and using different degrees of multiplexing. In most of these genomes, the expected motifs were unknown beforehand. Tables 4 (Ion Torrent) and 5 (Illumina MiSeq) show the number of unique reads from which each motif was derived, the total number of motif sites in the reference, and the fraction of all sites that was detected. Illumina reads ranged from 26-80 bases, and Ion Torrent reads from 20-80 bases. Reads of length 41-80 bases were used to determine background parameters, and those of lengths 26-40 bases were searched for motifs. As many as four motifs were discovered in a single genome by a single MFRE (*S. denitrificans* DSM1251 in Table 5).

**Table 5.**
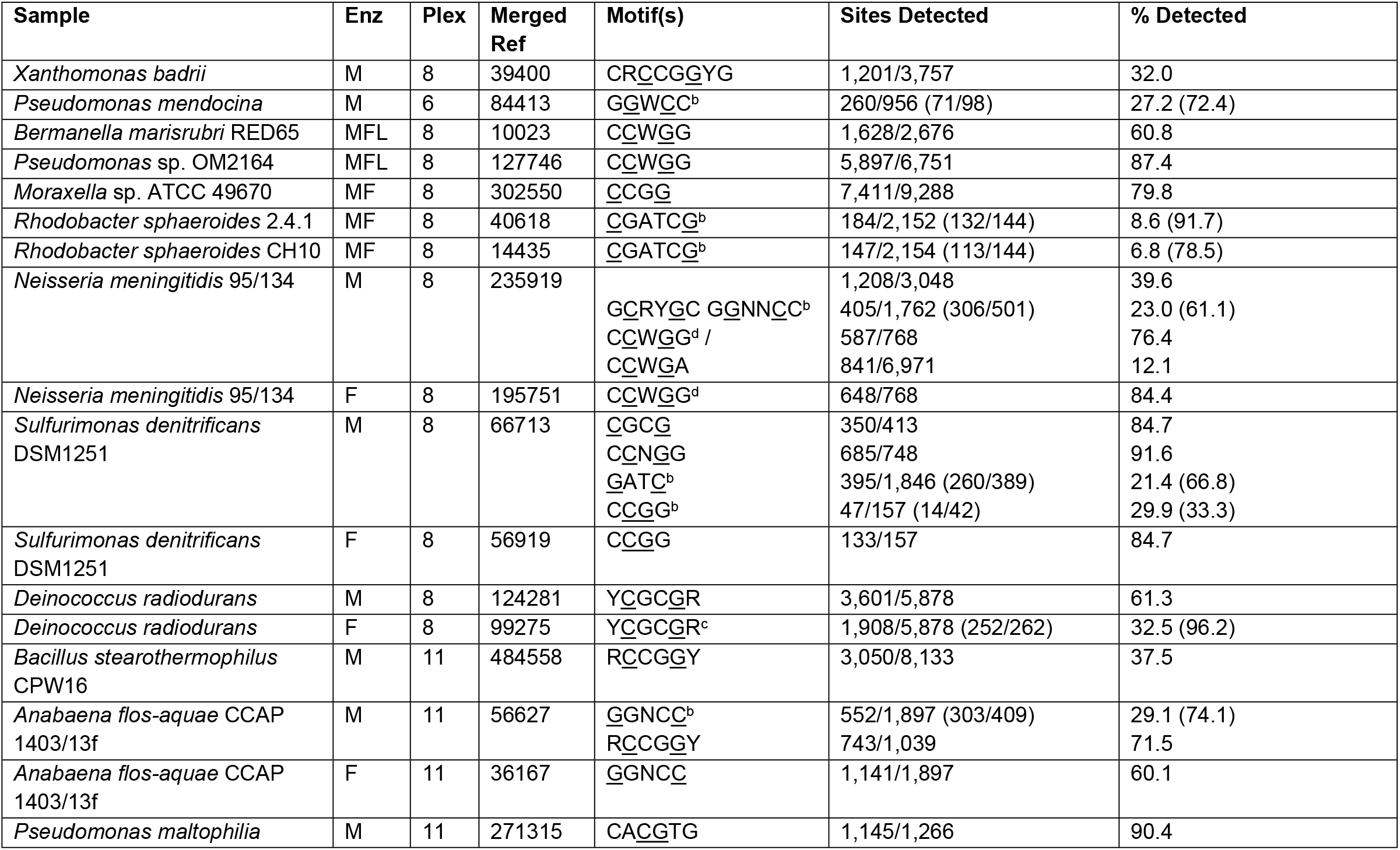
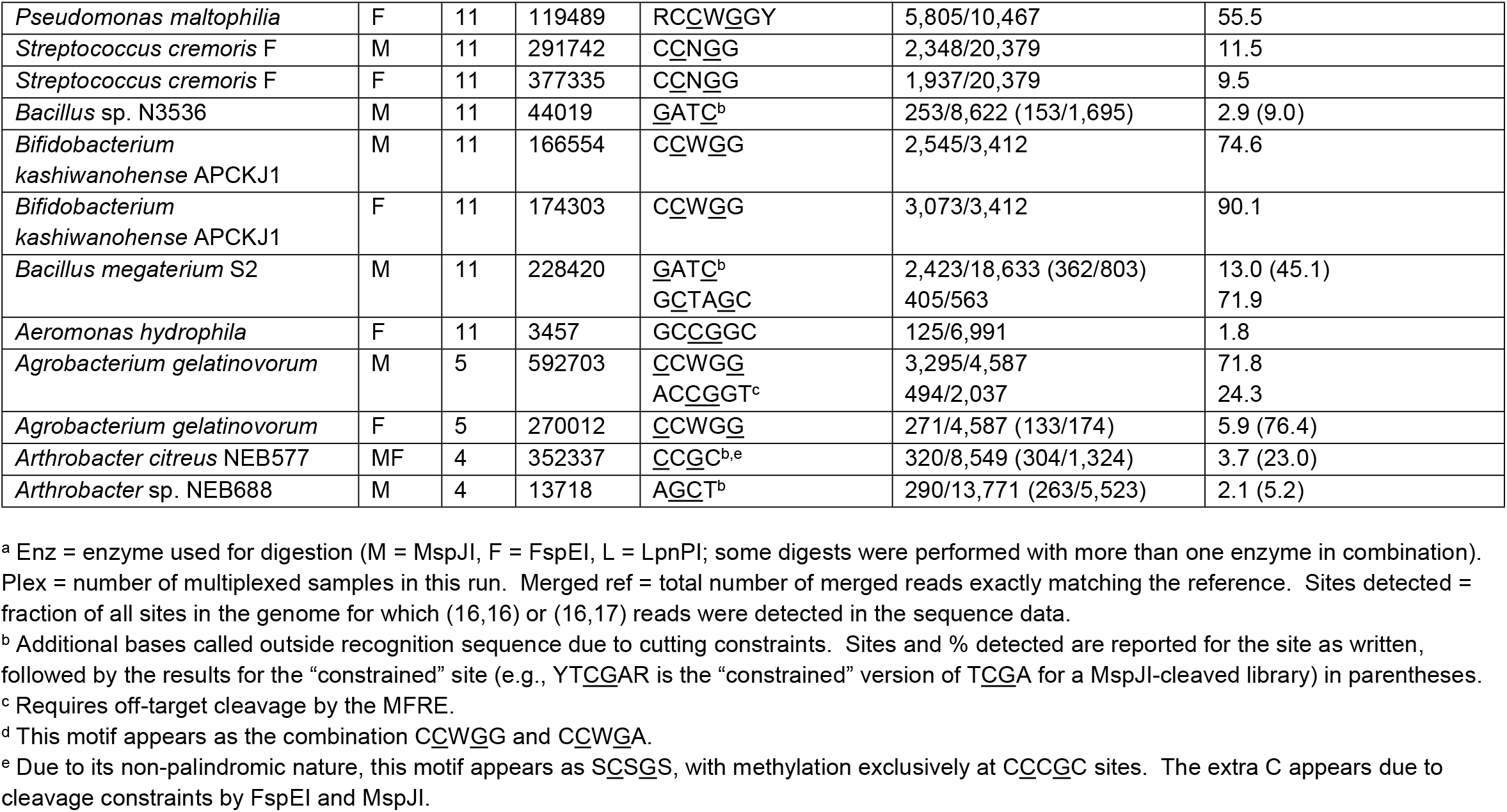
Motifs determined using the Ion Torrent platform.^a^

Motifs were successfully identified with as few as 51 unique reads, and from samples multiplexed to as many as 11 per run. In some cases, the MFRE’s own recognition site prevented cleavage of every instance of a MTase motif, and so the “apparent” motif (i.e., that determined automatically by the program) is over-constrained. For example, <u>G</u>AT<u>C</u> appears as *YNN*<u>G</u>AT<u>C</u>*NNR* when digested by MspJI, and <u>C</u>GATC<u>G</u> appears as *C*<u>C</u>GATC<u>G</u>*G* when digested by FspEI. (Bases in italics are part of the MFRE recognition site but not of the MTase motif.). These extraneous elements were removed manually. For these cases, the tables show data for both the “true” motif and the “apparent” (MFRE-cleavable) version of the motif, including the fraction of true and apparent sites represented in the sequence data. The difference between these two fractions was often large (see, for examples, <u>G</u>AT<u>C</u> and G<u>G</u>W<u>C</u>C in Table 4 and <u>C</u>GATC<u>G</u> in Tables 4 and 5).

In most cases, the deduced motif was derived from at least 30% of all motif sites in the genome, but in several cases the fraction of sites represented in the sequence data was lower, even when the site was not constrained by MFRE cleavage preferences. Even when the fraction of detected sites was low, we could often identify evidence that it was genuine. For example, AC<u>CG</u>GT was identified as a motif in *A. gelatinovorum* based on detection of only 24% of genomic sites in the sequence data (Table 5). This motif corresponds to that of AgeI, a REase previously characterized from this organism. In *S. cremoris* F, the site C<u>C</u>N<u>G</u>G was detected among only about 10% of sites by both MspJI and FspEI cleavage independently, but this activity (M.ScrFIA/B) has again been previously characterized (40). It appears in this case that a small number of M.ScrFI sites are highly overrepresented in the sequence data. And in *Arthrobacter* sp., the site A<u>GC</u>T was detected from only 5% of cleavable reads by MspJI. Although the fraction of sites is very low, the closest characterized homologs of the enzyme responsible, M.AscII, methylate this same site.

In certain cases, motifs were difficult to deduce due to significant off-target activity, presumably by the MTase. For example, in *Halorubrum* sp. BOL3-1 cleaved with MspJI, the apparent motif was BTCGAV (3265/33,268 = 9.8% sites represented). However, on closer inspection, this motif was composed of a canonical motif, CT<u>CG</u>AG (95.2% sites represented; Table 4), plus off-target activities at the asymmetric sites GT<u>C</u>GAG/CT<u>C</u>GAC (1937/12,621 = 15.3%) and TTCGAG/CTCGAA (780/5496 = 14.2%). Similarly, in *N. meningitidis* 95-134 cleaved with MspJI, the apparent motif YCWGR was composed of the canonical motif C<u>C</u>W<u>G</u>G (Table 5) plus off-target activity at the asymmetric site C<u>C</u>WGA/T<u>C</u>WGG (841/6971 = 12.1%).

A summary of the motifs identified in Tables 4 and 5 is shown in Table S4, and a summary of the genes responsible, arranged by genome, is shown in Table S5. In the 27 genomes under study (including the heterologous MTases expressed in two *E. coli* clones), 24 separate motifs were identified. The most common motifs found were C<u>C</u>W<u>G</u>G (5 genomes, not including the clones) and <u>G</u>AT<u>C</u> (4 genomes), with most motifs found in a single genome. All except one were palindromic. While this result may reflect inherent biases in the MFRE-Seq method (see Discussion), it does appear that the large majority of m5C motifs identified by other methods are also palindromic (Table S6). In the majority of the motifs we found (18 of 24), the top strand m5C was located 5’ to the bottom strand m5C, resulting in CCMD read lengths less than 33 bases. Motif lengths ranged from 4 to 8 bp and CCMD lengths ranged from 28 to 37 bases despite the fact that our search parameters permitted the identification of motifs and read lengths outside of these ranges. To date, no m5C MTases have been found to be associated with Type I or Type III R-M systems (16), and so all of the motifs found here likely belong to Type II R-M systems or to orphan MTases.

### Comparison with an independent method

As this manuscript was being prepared, a novel method for detecting m5C, EM-Seq, has been described (Vaisvila et al., manuscript submitted) and commercialized as a kit (New England Biolabs, Ipswich, MA). The kit relies on the same principle of C>U conversion as bisulfite sequencing but uses enzymatic rather than chemical methods to accomplish this. While MFRE-Seq is cheaper and less labor-intensive, we wished to validate our results with an independent method and used EM-Seq for this purpose. We compared results of MFRE-Seq and EM-Seq for two bacterial strains and three digests (*E. coli* DHB4 with MspJI, *E. coli* DHB4 with FspEI, and *P. mendocina* with MspJI) and found they yielded identical results (C<u>C</u>W<u>G</u>G in the case of both *E. coli* digests and G<u>G</u>W<u>C</u>C in the case of *P. mendocina*).

## DISCUSSION

Aside from the methylated base itself, the recognition site, or motif, is the primary distinguishing characteristic of bacterial DNA MTases. In the last ten years, the determination of MTase motifs has become commonplace due to the SMRT sequencing platform. However, results from SMRT sequencing have been uneven, with the vast majority of characterized examples being m6A and m4C MTases, for which the kinetic signals are pronounced. The study of m5C MTase has lagged behind due to the location of the methyl group. While methyl groups on m6A and m4C are directly involved with base pairing, that on m5C is not and is instead positioned in the major groove where it does not significantly contact the DNA polymerase (41), resulting in a more subtle perturbation of the kinetics of base incorporation. Roughly one third of all Type II R-M systems utilize m5C as the protective agent, and so alternative methods to characterize m5C MTase motifs are necessary to gain a complete picture of bacterial epigenetics.

In Type II R-M systems, the specificity determinants of paired MTases and REases are independent of each other. Because MTases were so rarely characterized prior to ten years ago, it was traditionally assumed that the recognition site of a MTase was identical to its cognate REase. SMRT sequencing results have shown that, for m6A and m4C MTases, this assumption holds true in most cases (see examples in REBASE). Using MFRE-Seq, we show here that it holds true of m5C MTases as well. Tables 4 and 5 include fourteen cases where an observed m5C activity can be matched with a characterized restriction enzyme from the same strain. In all cases, the MFRE-determined methylation motif matches exactly the recognition site of the known REase: HhaI, AvaII, PmeII, MspI, Rsp241I, SdeAII, BsrFI, AflI, PmlI, ScrFI, BscXII, BmtI, AgeI, and AciI. While it is reasonable to expect that the MTase of some Type II RM systems could have a broader specificity than the cognate REase, such cases appear to be relatively rare.

The above results notwithstanding, we observed several cases of apparent off-target activity, by either or both of the MTase and the MFRE. In all such cases we observed (notably in *Halorubrum* sp. BOL3-1 and *N. meningitidis* 95-134, discussed in Results), the off-target sites are asymmetric. For such sites to appear at appreciable frequency in the data, they must be cleaved on both sides of the site by the MFRE. This implies that (1) these sites are methylated on both strands, and (2) the sequences on both strands conform to the recognition site of the MFRE. Type II MTases typically act as monomers, methylating both strands independently. Asymmetric sequences typically require two MTases to achieve full methylation, so whether and how these off-target sequences are being methylated is at present unclear.

Furthermore, in the case of *Halorubrum* sp. BOL3-1, one of the asymmetric sites, GT<u>C</u>GAG/CT<u>C</u>GAC, should be cleavable by MspJI on only one side, even if methylated on both strands. The CT<u>C</u>GAC strand does not fit the <u>C</u>NNR pattern recognized by MspJI. We observed off-target cleavage by FspEI as well, in the case of *Deinococcus radiodurans*. MspJI cleavage discovers the motif Y<u>C</u>GC<u>G</u>R, with all four non-degenerate sequences represented in roughly equal fractions. Cleavage with FspEI should result in the apparent motif C<u>C</u>GC<u>G</u>G due to the MFRE’s cleavage requirements. However, we observed significant off-target activity at C<u>C</u>GCGA/T<u>C</u>GCGG sites (1,651/5,277 = 31.3%), one strand of which should not be cleavable. The nature of this activity is likewise unclear, but further examination may shed further insight into the cleavage requirements of MFREs. We are at present examining the phenomenon of “off-target” activity further.

Poor detection of sites by MFRE-Seq can be due either to low levels of methylation in the genome or to low numbers of sites. For example, the genome of *Anabaena variabilis* ATCC 27893 encodes four R-M systems with associated m5C MTases: M.AvaII (GGWCC), M.AvaIVP (predicted GCTNAGC, but possibly inactive), M.AvaVIII (CGATCG), and M.AvaIX (RCCGGY) (16). Using MFRE-Seq, we detected two of these motifs in the genomic DNA, R<u>C</u>CG<u>G</u>Y and <u>C</u>GATC<u>G</u>, but a third motif (G<u>G</u>W<u>C</u>C) was detectable only in an *E. coli* clone overexpressing M.AvaII, suggesting poor methylation in the native host. The <u>C</u>GATC<u>G</u> motif was detectable only with FspEI and manifested as VCGAYCGS, from which CGATCG was determined by parsing all 12 possible degenerate instances of this site (Table 7). The reason this site was so difficult to detect was that FspEI cleavage requirements restricted cutting primarily to CCGATCGG sites, for which there are only 4 in the entire genome (Table 5). The reason the motif was detected at all was due primarily to, again, off-target activity by FspEI at C<u>C</u>GATCGC/G<u>C</u>GATCGG sites (Table 7).

**Table 7.**
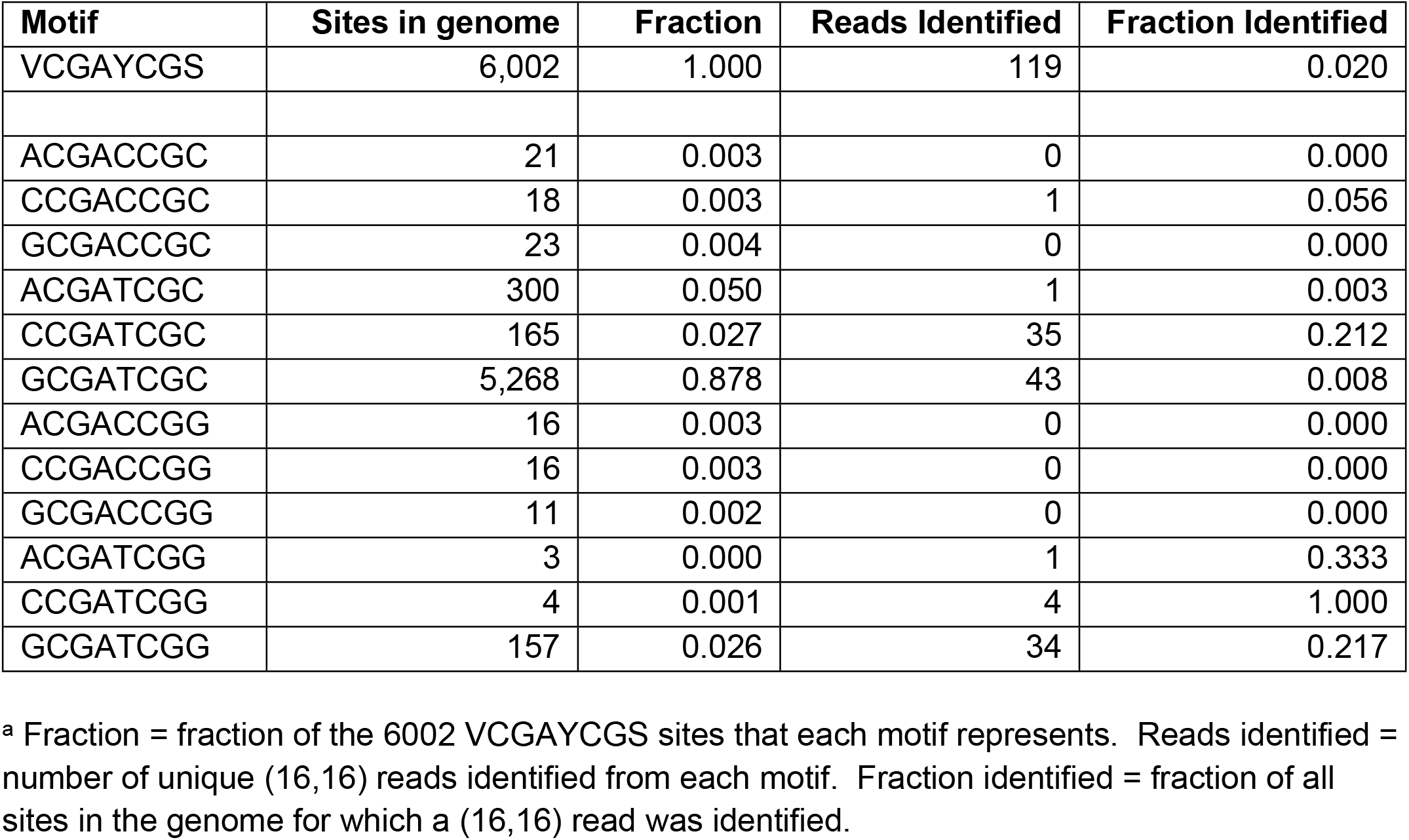
Read data for M.AvaVIII.^a^

| Motif | Sites in genome | Fraction | Reads Identified | Fraction Identified |
| --- | --- | --- | --- | --- |
| VCGAYCGS | 6,002 | 1.000 | 119 | 0.020 |
| ACGACCGC | 21 | 0.003 | 0 | 0.000 |
| CCGACCGC | 18 | 0.003 | 1 | 0.056 |
| GCGACCGC | 23 | 0.004 | 0 | 0.000 |
| ACGATCGC | 300 | 0.050 | 1 | 0.003 |
| CCGATCGC | 165 | 0.027 | 35 | 0.212 |
| GCGATCGC | 5,268 | 0.878 | 43 | 0.008 |
| ACGACCGG | 16 | 0.003 | 0 | 0.000 |
| CCGACCGG | 16 | 0.003 | 0 | 0.000 |
| GCGACCGG | 11 | 0.002 | 0 | 0.000 |
| ACGATCGG | 3 | 0.000 | 1 | 0.333 |
| CCGATCGG | 4 | 0.001 | 4 | 1.000 |
| GCGATCGG | 157 | 0.026 | 34 | 0.217 |

<sup>a</sup> Fraction = fraction of the 6002 VCGAYCGS sites that each motif represents. Reads identified = number of unique (16,16) reads identified from each motif. Fraction identified = fraction of all sites in the genome for which a (16,16) read was identified.

The accuracy of MFRE-Seq depends on the availability of a set of methylated examples that is both sufficiently large and unbiased. Our randomization tests show that, at the level of “non-CCMD reads” we typically see, on the order of 50 examples or fewer are needed. In an unbiased sequence, a fully specified 8 bp motif should occur about 50 times in a 3.3 Mbp genome, meaning even the longest known motifs should be detectable by MFRE-Seq. That said, genomes are not random sequences, but can have significant biases for or against specific *k*-mers. The case of A. variabilis mentioned above is an extreme example: there are 5,268 GCGATCGC sites, but only 4 CCGATCGG sites. All methods of determining sequence motifs suffer equally from this same difficulty. Although this challenge can be overcome by testing batteries of equally frequent sites, this type of experiment is beyond the scope of this work.

The method we have described here, which we term MFRE-Seq, has both advantages and disadvantages relative to other methods for motif determination. The primary advantage is ease of use, in that it requires only REase digestion of the DNA sample prior to library preparation, as opposed to damaging sodium bisulfite chemical treatment. It is compatible with both Ion Torrent and Illumina sequencing platforms; SMRT sequencing of the fragments is not recommended due to the very short nature of the MFRE-derived library inserts. Processing of sequence data to derive motifs is also straightforward. We have presented one possible method, namely identifying CCMD reads and deriving the motif by simple alignment. However, other methods could easily be used instead, including searching for overrepresented sequences using MEME (32) or Mosdi (42), building motifs from probable m5C sites using MotifMaker, and other methods.

We have used this method in conjunction with a reference sequence, using exact matching of sequence reads as a filtering step to remove reads with potential errors and reads derived from non-reference sources. After reducing sequence reads derived from unproductive sources such as the activator oligonucleotide and molecular weight markers (see Results), the fraction of reads not exactly matching the reference has tended to be low. In our experiments with *E. coli* DHB4, performed after these steps were implemented, more than 96% of reads were exact matches to the reference (Table 1), so it may also be possible to use MFRE-Seq in the absence of a reference, perhaps using read copy number as a filtering step.

While MFRE-Seq has important advantages over bisulfite-based methods, there are certain types of methylated sites that are either not straightforward or impossible for it to identify. Sites that are not straightforward are nonpalindromic sites and hemi-methylated sites. Nonpalindromic sites that are cleavable on both strands by one or more MFREs will, using the motif-determination method described here, generate degenerate motifs that represent the average of the motif sequences on the two strands. Deconvolution of this data by examining the representation of each non-degenerate instance will make the true site apparent, and we did this successfully in the case of AciI (methylated at <u>C</u>C<u>G</u>C). Hemi-methylated sites are in theory discoverable, but since they are cut by the MFRE on only one side, an alternative motif-searching method must be used. It should be noted that some R-M systems rely on two separate MTases to methylate both strands of an asymmetric site. Those cases where one of those MTases is either m6A or m4C are, for the purposes of MFRE-Seq, considered hemi-methylated, as they are methylated at m5C on only one strand.

Those methylated sites (whether palindromic, nonpalindromic, or hemi-methylated) that are at present impossible to identify are those that do not conform to the recognition sites of the available MFREs, which as of this writing comprises MspJI, FspEI, and LpnPI. (We did nonetheless identify a small number of such sites in our data, presumably due to off-target activity by the MFREs, but such cases appear to be rare.) Table S6 shows all known m5C DNA MTase recognition sites found in REBASE and their MFRE cleavage properties. There are 100 different sites, of which 72 are palindromic and 82 are m5C-methylated on both strands, making them discoverable with MFRE-Seq using the computational method described here. 68 of the 82 are cleavable with either MspJI or FspEI. However, in many cases only a subset of instances of the site can be cleaved due to the MFRE’s own recognition properties. For examples, <u>G</u>AT<u>C</u> sites are only cleavable by MspJI when they fit the profile YNN<u>G</u>AT<u>C</u>NNR, and <u>C</u>AT<u>G</u> sites are only cleavable by FspEI when they fit the profile C<u>C</u>AT<u>G</u>G. MFRE recognition sites need to be taken into account to avoid over-specification of MTase motifs.

Identification and characterization of additional MFRE family members with orthogonal specificities should help overcome this limitation, and this work is currently in progress. In the meantime, MFRE-Seq has already identified numerous new recognition sites and methylated bases within known sites, and it serves as a useful complement to other methods of m5C motif determination.

## ACKNOWLEDGMENTS

We thank Mehmet Berkmen, Francesca Bottacini, Priya and Shil DasSarma, Christopher D. Johnston, Katherine P. Lemon, Richard D. Morgan, Bianca Stenmark, and Jill Zeilstra for providing strains, genomic DNA, and/or clones for analysis. The authors would also like to thank Don Comb for support.

## SUPPORTING INFORMATION

Text S1. Description of Data Sets S1 through S4.

Table S1. Base filtering of selected read structures containing C<u>C</u>W<u>G</u>G (*l*_–2_ = 31) or <u>C</u>CWG<u>G</u> (*l*_–4_ = 29).

Table S2. Number of (all / base-filtered) reads of various classes for lengths 30-32 bases. Table S3. Reads from enzyme activator oligonucleotides.

Table S4. Summary of m5C motifs identified in Tables 4 and 5.

Table S5. REBASE nomenclature for motifs discovered or confirmed with MFRE-Seq. Table S6. MFRE cleavage properties of all known m5C motifs.

Figure S1. Venn diagram of unrepresented sites from the four libraries of *E. coli* K-12 DHB4 described in Table 1 (with corresponding numbering). Figures show the number of loci common to one or more libraries.

Figure S2. Gel photo (20% TBE-PAGE stained with SYBR Gold) of MFRE digest reactions showing activator and cleavage bands. All reactions were MspJI (1 µL) digests of *E. coli* DHB4 genomic DNA with 0.525 µM enzyme activator in 40 µL 1x CutSmart buffer, 37°C 3 hrs. M = Low Molecular Weight DNA Ladder (New England Biolabs). Lanes 1-5 contained 3 µg genomic DNA, and lanes 6-10 contained 1.5 µg genomic DNA. Activators: none (lanes 1 and 6), standard activator (lanes 2 and 7), activator-N (lanes 3 and 8), activator-U (lanes 4 and 9), or activator-UN (lanes 5 and 10).

Figure S3. Examples of read distribution with 3 digest cleanup protocols. All 3 samples were digested with MspJI and sequenced on the Ion Torrent platform. A. One-step spin-column cleanup, which keeps all fragments, small and large; *Arthrobacter* sp., CCMD length = 34. B. Two-step spin-column cleanup, which selects for fragments < 100 bp; *E. coli* DHB4, CCMD length = 31. C. Gel-purification of small fragments (20-50 bp range) from 20% polyacrylamide; *E. coli* DHB4, CCMD length = 31. Data Sets S1-S4. Compressed archives of processed MFRE-Seq FASTA files from which motifs in Tables 4 and 5 were derived.

